# Functional oscillation of a multienzyme glucosome assembly during cell cycle progression

**DOI:** 10.1101/2022.01.06.475270

**Authors:** Miji Jeon, Danielle L. Schmitt, Minjoung Kyoung, Songon An

## Abstract

Glucose metabolism has been studied extensively to understand functional interplays between metabolism and a cell cycle. However, our understanding of cell cycle-dependent metabolic adaptation particularly in human cells remains largely elusive. Meanwhile, human enzymes in glucose metabolism are shown to functionally organize into three different sizes of a multienzyme metabolic assembly, the glucosome, to regulate glucose flux in a size-dependent manner. Here, using fluorescence single-cell imaging techniques, we discover that glucosomes spatiotemporally oscillate during a cell cycle in an assembly size-dependent manner. Importantly, their oscillation at single-cell levels is in accordance with functional contributions of glucose metabolism to cell cycle progression at a population level. Collectively, we demonstrate functional oscillation of glucosomes during cell cycle progression and thus their biological significance to human cell biology.

## Introduction

Metabolism undergoes dynamic changes during a cell cycle to provide energy and building blocks for a cell to duplicate and divide (*1*–*4*). Glucose metabolism, particularly including glycolysis and the pentose phosphate pathway, are upregulated during the G1 phase. Their metabolic activities remain high during the S phase to support DNA replication. However, when DNA replication is completed in the G2 phase, glucose metabolism decreases to their basal levels. Subsequently, when a cell enters the G2 phase, glycolysis remains at the basal level but protein and lipid synthesis is promoted during the G2 phase. Then, during the mitosis, a cell divides into two daughter cells. The life cycle of eukaryotic cells continues along with described metabolic oscillation when daughter cells reenter a cell cycle. However, other than in yeast and unicellular eukaryotic cells, our understanding of such cell cycle-dependent metabolic demands particularly in human cells has been largely elusive (*5*, *6*).

Nevertheless, mounting evidence in recent years has started suggesting functional interplays between glucose metabolism and a cell cycle in human cells (*4*–*6*). For instance, expression of hexokinase 2 in human epithelial type 2 cells and cancer-associated fibroblasts was upregulated during the G1 phase, indicating the promotion of glycolysis (*7*, *8*). The activity of 6-phosphofructo-2-kinase/fructose-1,6-bisphosphatase isoform 3 (PFKFB3), which allosterically activates phosphofructokinase 1 (PFK) in glycolysis, was also increased in the late G1 but decreased in the early S phase in human embryonic kidney cells, human breast cancer cells (MDA-MB-231 and MCF7), neuroblastoma cells (SHSY-5Y) and human cervical cancer cells (HeLa) (*9*–*12*). In addition, enzymatic activity of glucose-6-phosphate dehydrogenase in the pentose phosphate pathway was enhanced during the late G1 and the early S phase of a cell cycle in human colon adenocarcinoma cells (HT29) (*13*, *14*). While metabolic interplays with a cell cycle in human cells are indeed gradually disclosed, however, these studies and others (*4*–*6*) have mainly tracked the activity change of a representative enzyme in given metabolic pathways to deduce a relationship between metabolism and a cell cycle. Therefore, it has been barely demonstrated yet how metabolic pathways themselves oscillate to functionally contribute to a cell cycle in human cells.

Meanwhile, we have identified that human liver-type phosphofructokinase 1 (PFKL) spatially forms various sizes of a cytoplasmic metabolic assembly, namely the glucosome, and recruits at least three rate-determining enzymes in glucose metabolism, including fructose-1,6-bisphosphatase, pyruvate kinase, and phosphoenolpyruvate carboxykinase 1 (*15*). Due to the heterogeneity of the observed sizes of the glucosomes under fluorescence live-cell microscopy, we have subsequently categorized them into three subclasses for their functional characterization (i.e., small-, medium-, and large-sized glucosomes) (*15*). Briefly, small-sized glucosomes are defined as having less than our calculated area of a point spread function for the emission of monomeric enhanced green fluorescent protein (mEGFP) (i.e., ~ 0.1 μm^2^). Medium-sized glucosomes are defined to have larger than 0.1 μm^2^ but less than 3 μm^2^ in size. The 3 μm^2^ cutoff is experimentally determined based on the fact that non-cancerous human cell lines we have tested do not display glucosomes that are larger than 3 μm^2^ in our growth conditions. Accordingly, large-sized glucosomes are defined to range from 3 μm^2^ to 8 μm^2^, which have been detected in various human cancer cell lines (*15*). Nevertheless, when fluorescent granules display larger than 8 μm_2_ in size at single-cell levels, we have found that such aggregations are primarily composed of an immobile fraction of transfected enzymes and importantly their formation is apparently non-specific to the identity of transfected enzymes (*15*, *16*). In this work, fluorescent aggregates larger than 8 μm^2^ in size are not classified as glucosomes. Others have also detected various sizes of spatial assemblies of human PFKL, for example, in HepG2 cells by immunostaining (*17*) as well as in rat breast cancer cells by fluorescence live-cell imaging (*18*). Most importantly, our localization-function studies using high-content live-cell imaging assays and mathematical modeling analysis (*15*, *19*) have strongly suggested that small-sized glucosomes primarily promote glycolysis whereas medium- and large-sized glucosomes shunt glucose flux preferentially into the pentose phosphate pathway and serine biosynthesis, respectively. Therefore, we envision that glucosomes would be functionally associated with various biological processes in human cells.

In this work, we have investigated metabolic contributions of glucosomes to cell cycle progression by employing fluorescence live-cell imaging along with cell synchronization and flow cytometry. Our comprehensive analysis has now revealed that small-sized glucosomes drastically oscillate along with a cell cycle and medium-sized glucosomes are upregulated only during the G1 phase whereas large-sized glucosomes barely or minimally oscillate during a cell cycle. Importantly, should we consider our localization-function studies that show the assembly size-dependent functional roles of glucosomes (*15*, *19*), our results here are indeed consistent with current understanding of metabolic demands at the G1 phase (i.e., upregulation of glycolysis and the pentose phosphate pathway) and further provide heretofore unrecognized metabolic demands at the G2 phase, which involve the downregulation of glycolysis but relatively sustained glucose flux into the pentose phosphate pathway and serine biosynthesis. Collectively, we provide compelling evidence that glucosomes dynamically oscillate during cell cycle progression in a size-dependent manner to regulate glucose flux between glycolysis and building block biosynthesis, thereby demonstrating functional significance of glucosomes and their dynamics in living human cells.

## Results

### Expression of an intracellular glucosome marker for cell cycle analysis

Previously, PFKL has been implicated as a scaffolder of a multienzyme metabolic assembly for glucose metabolism in eukaryotic cells (*15*, *17*, *18*). Of particular, human PFKL with a fluorescent protein tag (e.g., PFKL-mEGFP) has been used as an intracellular marker for human multienzyme assembly, the glucosome, in living human cells (*15*, *19*). In this work, we first carried out flow cytometry-assisted cell cycle analysis with asynchronized human triplenegative breast carcinoma Hs578T cells with and without the expression of PFKL-mEGFP. Our histogram analysis of asynchronous Hs578T cells showed that the percentage of cells in each phase of a cell cycle was statistically unchanged in the presence of PFKL-mEGFP (**Fig. 1**). Apparently, the expression of PFKL-mEGFP didn’t alter or negatively influence the overall distribution of Hs578T cells among the phases of a cell cycle. Therefore, we determined that subcellular dynamics of PFKL-mEGFP in living human cells would be adequate as an intracellular glucosome marker to find out a functional interplay between glucose metabolism and cell cycle progression.

**Fig. 1.**
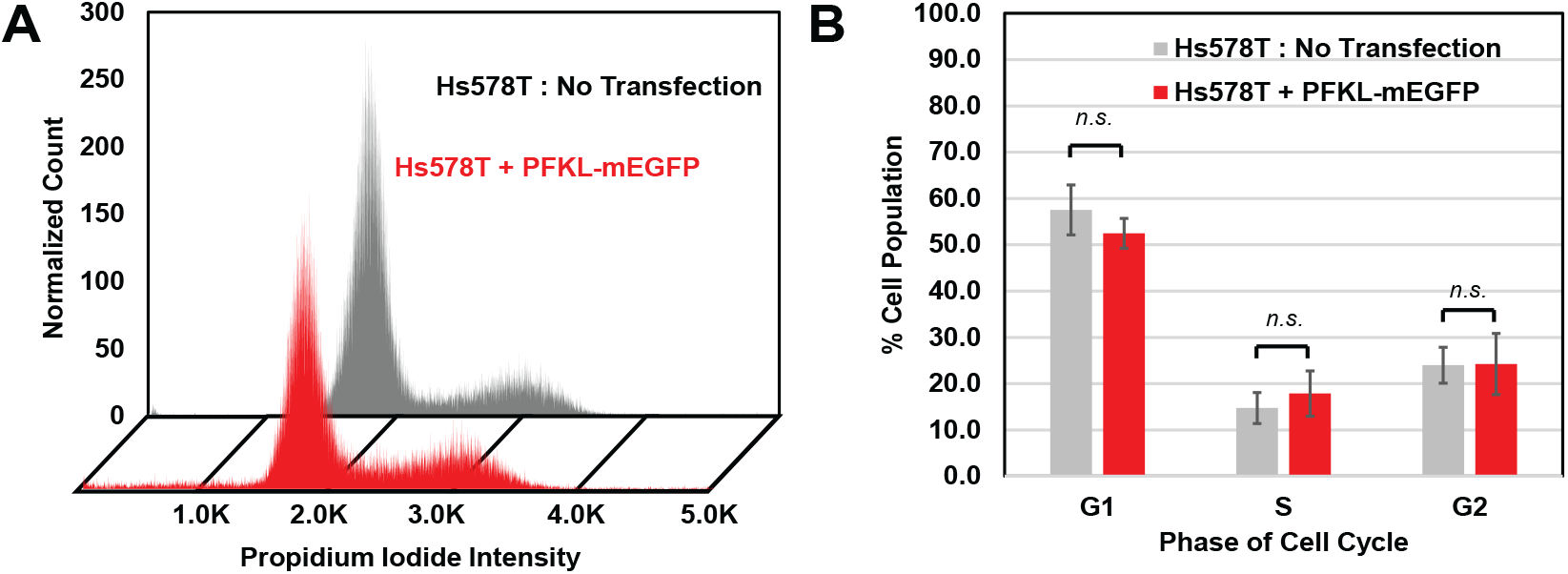
Expression of a glucosome marker, PFKL-mEGFP, in Hs578T cells. **(A)** A cell cycle analysis by flow cytometry showed the distribution of Hs578T cells with (red) and without (light grey) expressing PFKL-mEGFP. **(B)** The percentages of cell population in each phase of a cell cycle were graphed with (red, *N* = 9) and without (light grey, *N* = 6) the expression of PFKL-mEGFP in Hs578T cells. Statistical analyses were performed using two-way ANOVA and Tukey’s multiple comparison tests. Error bars represents standard deviations that are obtained from six to nine independent trials. Statistical significance is defined as *p* < 0.05 with a 95% confidence interval while n.s. refers to not significant.

### Small molecule-promoted cell synchronization with a glucosome marker

We then investigated cell cycle-dependent distribution of synchronized Hs578T cells with and without a glucosome marker, PFKL-mEGFP (**Fig. 2A**). First, we treated Hs578T cells with lovastatin (40 μM), which is known to arrest cells in the G1 phase (*20*, *21*). Without glucosomes being expressed, the percentage of G1-arrested Hs578T cells increased ~16.6 % in the presence of lovastatin, relative to the percentage of asynchronous Hs578T cells in the G1 phase (i.e., 57.6 % to 74.2 %) (**Fig. 2B**, light grey vs. blue). Meanwhile, lovastatin also increased ~17.9 % of glucosome-transfected Hs578T cells in the G1 phase, relative to the percentage of asynchronous Hs578T cells expressing glucosomes, (i.e., 52.5 % to 70.4 %) (**Fig. 2C**, red vs. blue). This data supports that the expression of a glucosome marker has no significant influence on arresting cells in the G1 phase by lovastatin (**Fig. 2B** vs **2C**, blue).

**Fig. 2.**
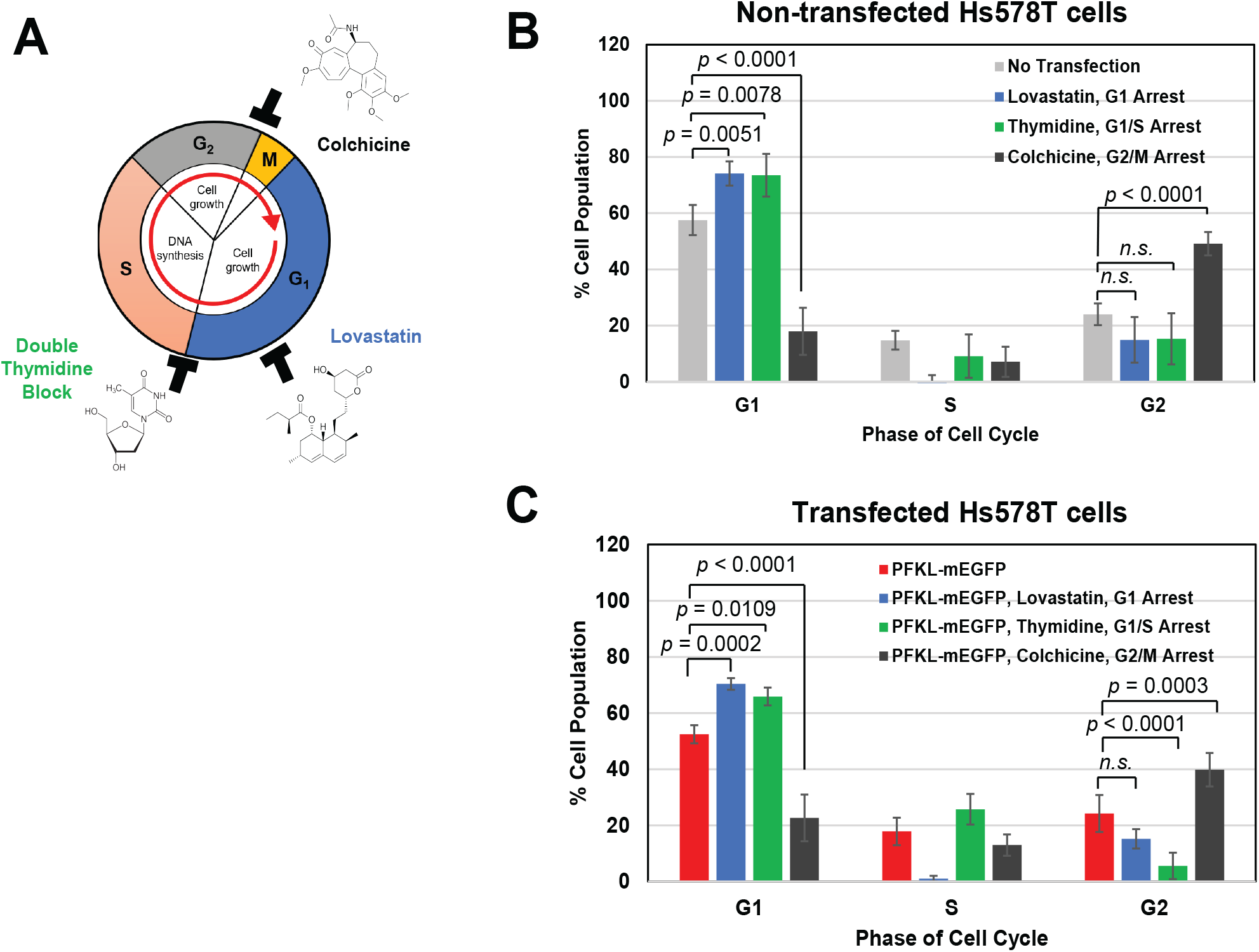
Small molecule-promoted cell synchronization with and without the expression of glucosomes in Hs578T cells. **(A)** A schematic diagram shows how small molecule-promoted cell synchronization is achieved based on their mechanisms of action at a given specific phase of a cell cycle. **(B-C)** Population changes of Hs578T cells at each phase of a cell cycle were graphed in the presence of a cell cycle inhibitor (i.e., lovastatin (blue), thymidine (green) and colchicine (black)) without (B, *N_lovastatin_* = 4, *N_thymidine_* = 4, *N_colchine_* = 4) and with (C, *N_lovastatin_* = 3, *N_thymidine_* = 3, *N_colchicine_* = 4) the expression of PFKL-mEGFP as a glucosome marker in Hs578T cells. Statistical analyses were performed using two-way ANOVA and Tukey’s multiple comparison tests. Error bars represents standard deviations that are obtained from three to nine independent trials. Statistical significance is defined as *p* < 0.05 with a 95% confidence interval while n.s. refers to not significant.

Second, we also used a double thymidine block (2 mM) to arrest Hs578T cells in the G1/S phase transition. Thymidine is capable of arresting cells at the G1/S boundary by blocking DNA replication (*22*, *23*). The treatment of thymidine showed a ~16.0 % increase of nontransfected Hs578T cells in the G1 phase relative to the G1 population of asynchronous nontransfected Hs578T cells (i.e., 57.6 % to 73.6 %) (**Fig. 2B**, light grey vs. green). Similarly, the percentage of glucosome-expressing Hs578T cells in the G1 phase was also increased by ~13.4 %, compared to the G1 population of asynchronous but glucosome-expressing cells (i.e., 52.5 % to 65.9 %) (**Fig. 2C**, red vs. green). As anticipated, the treatment of thymidine showed a similar trend as lovastatin did with comparable degrees of increasing the G1 population regardless of the glucosome expression (blue and green in **Fig. 2B** vs. **2C**).

Third, we blocked cell cycle progression in the G2/M phase using colchicine (100 nM), which is known to disrupt an interaction between alpha- and beta-tubulin for microtubule polymerization and thus interrupt the G2/M transition (*24*, *25*). Our data showed that in the absence of glucosomes, the percentage of Hs578T cells in the G2 phase was increased by ~25.1 % relative to the percentage of asynchronous Hs578T cells in the G2 phase (i.e., 24.0 % to 49.1 %) (**Fig. 2B**, light grey vs. black). In the presence of glucosomes, the percentage of Hs578T cells in the G2 phase was also increased ~15.6 % relative to the G2-arrested percentage of asynchronous Hs578T cells with glucosomes (i.e., 24.2 % to 39.8 %) (**Fig. 2C**, red vs. black). Indeed, regardless of the glucosome expression, a significant percentage of Hs578T cells was arrested in the G2 phase by the treatment of colchicine.

Taken all together, we demonstrated that Hs578T cells that express PFKL-mEGFP as a glucosome marker went through a cell cycle in a very similar way as non-transfected Hs578T cells did. Therefore, these results provided a solid baseline of the cell cycle-dependent distribution of glucosome-expressing Hs578T cells (**Fig. 1B**, red, or **Fig. 2C**, red) for our indepth cell cycle analysis.

### Impacts of cell cycle inhibitors on cell population showing differently sized glucosomes

Now, to understand the impacts of a cell cycle inhibitor on glucosome assemblies in Hs578T cells, we analyzed the percentage of Hs578T cells displaying different sizes of glucosomes in the absence and presence of each cell cycle inhibitor. First, our high-content imaging analysis with lovastatin (20 μM) revealed that the population of Hs578T cells showing large-sized glucosomes drastically decreased from 26.5 % to 12.0 % (*p* < 0.0001) (**Fig. 3A** and **Supplementary Fig. S1**). The population of cells showing small- and medium-sized glucosomes concurrently increased after the treatment of lovastatin; 58.8 % to 66.9 % (*p* < 0.05) and 13.3 % to 21.1 % (*p* < 0.05), respectively. In addition, when colchicine (10 nM) was added to arrest Hs578T cells in the G2 phase, the population of Hs578T cells showing large-sized glucosomes significantly decreased again from 26.5 % to 13.6 % (*p* = 0.0002) (**Fig. 3B** and **Supplementary Fig. S2**). Accordingly, the population of Hs578T cells showing small-sized glucosomes was significantly increased from 58.8 % to 75.1 % by colchicine (*p* < 0.0001). However, no change in the population of cells showing medium-sized glucosomes was observed (i.e., 13.3 % vs. 11.4 %, *p* > 0.05). These data revealed that colchicine modulated glucosome dynamics similarly at single-cell levels as lovastatin did, but colchicine impacted differently from lovastatin on the population of cells particularly that displayed medium-sized glucosomes. Considering that changes of cell population showing different sizes of glucosomes are indicative of metabolic alterations of the cells (*15*, *19*), our results suggest that collective influence of a cell cycle inhibitor on glucose metabolism at a population level appears to be different from its influences on glucosome dynamics at single-cell levels.

**Fig. 3.**
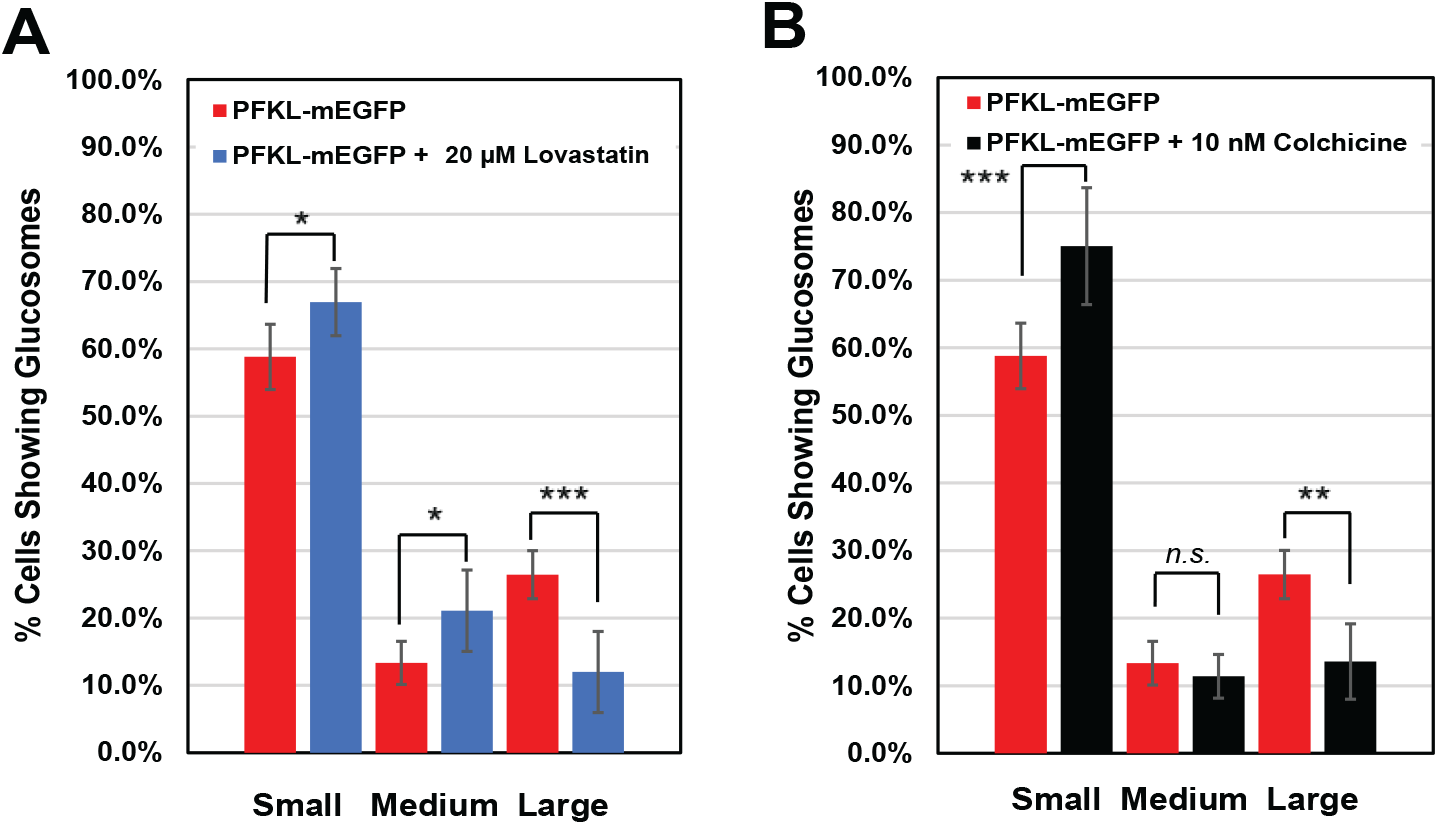
Effects of lovastatin and colchicine on cell population showing differently sized glucosomes. The population (%) of Hs578T cells displaying each size of PFKL-mEGFP assemblies was analyzed in the absence (*N* = 10) and presence (*N_lovastatin_* = 5 and *N_colchicine_* = 5) of 20 μM lovastatin **(A)** and 10 nM colchicine **(B)**, respectively. Statistical analyses were performed using two-way ANOVA and Tukey’s multiple comparison tests. Error bars represent standard deviations from at least five independent trials. Statistical significance is defined as *p* < 0.05 with a 95% confidence interval. * *p* < 0.05, \*\**p* < 0.001, *** *p* < 0.0001. n.s.; not significant.

### Size-dependent distribution of glucosomes at specific phases of a cell cycle in single cells

Next, to unambiguously examine whether or not a specific size of glucosomes is preferentially utilized at a specific phase of a cell cycle at single-cell levels, we introduced a cell cycle indicator, Vybrant^®^ DyeCycle^™^ Orange Stain (0.5 μM) (*26*, *27*), to Hs578T cells that expressed PFKL-mEGFP as a glucosome marker (**Fig. 4A**). We then randomly selected glucosome-positive Hs578T cells in the presence of lovastatin and colchicine, respectively. We subsequently quantified fluorescent intensities of the cell cycle indicator from the nucleus of the selected cells to determine at which phase of a cell cycle they were. Accordingly, a histogram was then set for a cell cycle analysis (**Fig. 4B**) as described in the Method section. As a control, we also analyzed glucosome-positive cells that were stained with the cell cycle indicator in the same way after the treatment of a vehicle, water (*N_vehicle_* = 172).

**Fig. 4.**
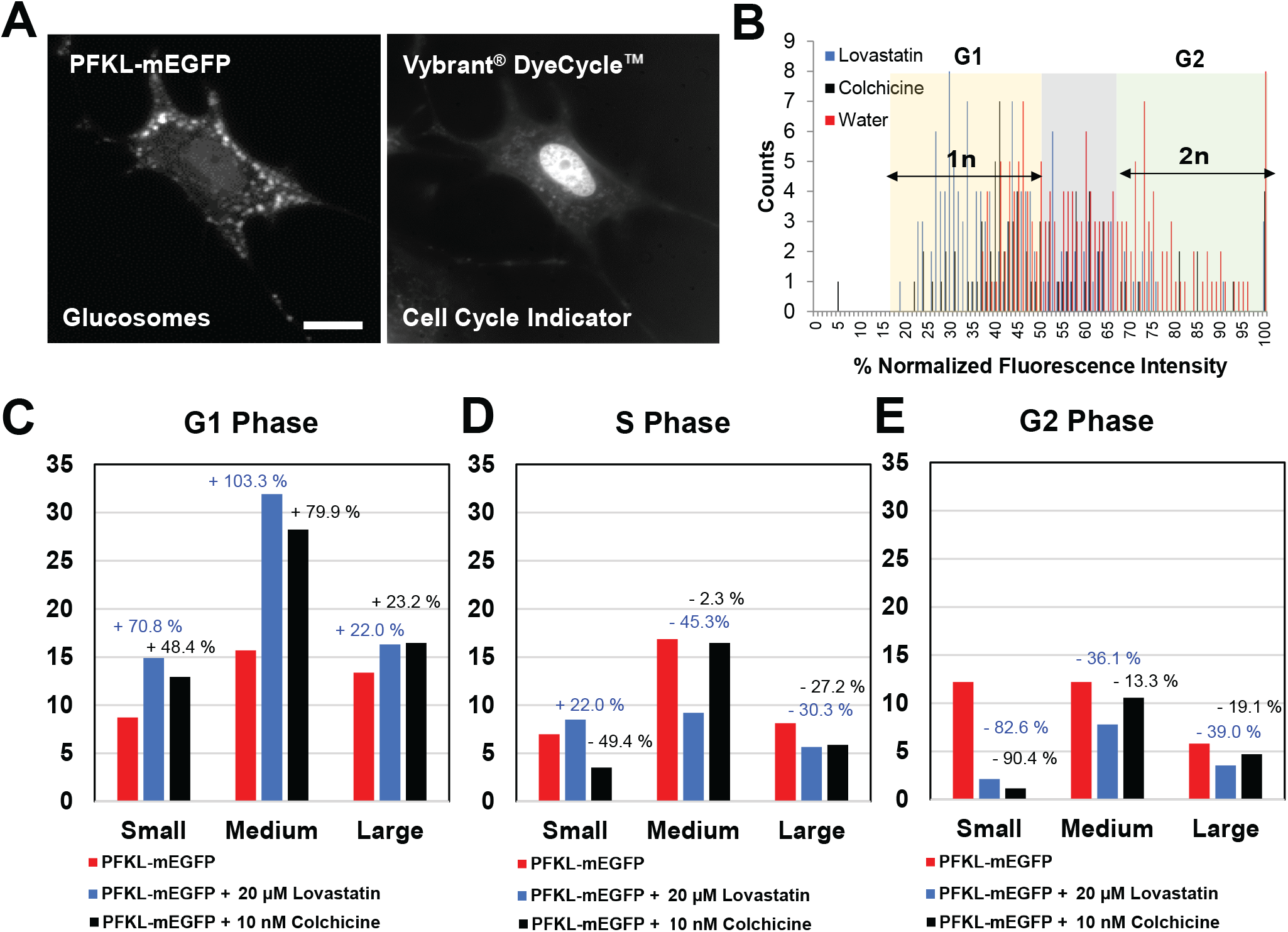
Glucosome dynamics during cell cycle progression in single cells. **(A)** A representative Hs578T cell expressing PFKL-mEGFP in the cytoplasm was stained with a cell cycle indicator, Vybrant^®^ DyeCycle^™^ Orange Stain, that localized in the nucleus. **(B)** A total of 398 cells (*N_lovastatin_* = 141, *N_colchicine_* = 85, and *N_vehicle_* = 172) that displayed both glucosomes and the cell cycle indicator were randomly selected from at least 9 independent trials. A histogram of the selected cells was then graphed the number of counted cells over the normalized fluorescence intensity. Accordingly, the 2n group represents Hs578T cells in the G2 phase while the 1n group represents Hs578T cells in the G1 phase. **(C-E)** Changes of the number of Hs578T cells displaying each size of glucosomes were analyzed in the presence of either lovastatin (blue) or colchicine (black) at each phase of the cell cycle. Percentage (%) changes of the number of single cells showing different sizes of glucosomes are annotated relative to a control (red).

When we analyzed the randomly selected Hs578T cells after the treatment of lovastatin (*N*_lovastatin_ = 141) and colchicine (*N*_colchicine_ = 85), respectively, we found that the number of G1-arrested Hs578T cells displaying all three sizes of glucosomes were increased relative to the controls (**Fig. 4C**, from red to blue or black). Although one specific size of glucosomes wasn’t distinctively regulated in the G1 phase, the number of Hs578T cells showing small- and medium-sized glucosomes in the G1 phase were drastically increased by lovastatin as well as by colchicine (**Fig. 4C**, red to blue or black). However, the numbers of Hs578T cells in the G2 phase, regardless of glucosome sizes, were decreased in the presence of lovastatin and colchicine, respectively, (**Fig. 4E**, red to blue or black). Particularly, we observed significant reduction of the number of G2-arrested Hs578T cells when they displayed small-sized glucosomes prior to the inhibitor treatment. We also noticed that the up- and down-regulation trends we observed with lovastatin at single-cell levels (**Fig. 4C-E**, red to blue) were very similar with the trends we observed with colchicine (**Fig. 4C-E**, red to black), except for cells in the S phase with small- and medium-sized glucosomes. Our data indicates that the mechanisms or consequences of action of lovastatin and colchicine don’t govern the utilization of specific sizes of glucosomes at single-cell levels. Instead, glucosome dynamics at single-cell levels appears to depend on at which phase of a cell cycle a cell is being during cell cycle progression. Collectively, we demonstrate that human cells are capable of dynamically regulating various sizes of glucosomes at subcellular levels during a cell cycle.

### Functional oscillation of glucosomes during cell cycle progression

To advance our understanding of the size-dependent dynamics of glucosomes during cell cycle progression, we have further analyzed our data (**Fig. 4**) showing the changes of glucosome sizes at each phase of a cell cycle. Regardless of the treatment of lovastatin or colchicine, we found consistently that on average the number of cells showing small-sized glucosomes in the G1 phase increased ~ 60 % while the number of cells showing small-sized glucosomes in the G2 phase significantly decreased ~ 86 % (**Fig. 5A**). This suggests that small-sized glucosomes are most dynamically associated with a cell cycle, indicating their oscillatory behavior during a cell cycle. Meanwhile, medium-sized glucosomes were upregulated only in the G1 phase by showing on average ~ 90 % increase of the number of cells (**Fig. 5B**), which suggests the importance of their metabolic contribution to the G1 phase. Additionally, large-sized glucosomes were barely or minimally associated with cell cycle progression (**Fig. 5C**), indicating their contribution remains relatively constant throughout a cell cycle. Finally, when we put all the data together in multiple cycles, our results clearly revealed that glucosomes oscillated along with a cell cycle in an assembly size-dependent manner (**Fig. 5D**). Considering that glucosome assemblies are functionally active in regulation of glucose flux between glycolysis and building block biosynthesis (*15*, *19*) (**Fig. 6A**), our work here demonstrate the size-dependent functional oscillation of glucosomes during cell cycle progression (**Fig. 6B**), thereby revealing metabolic pathway-level functional interplays between glucose metabolism and a cell cycle in human cells.

**Fig. 5.**
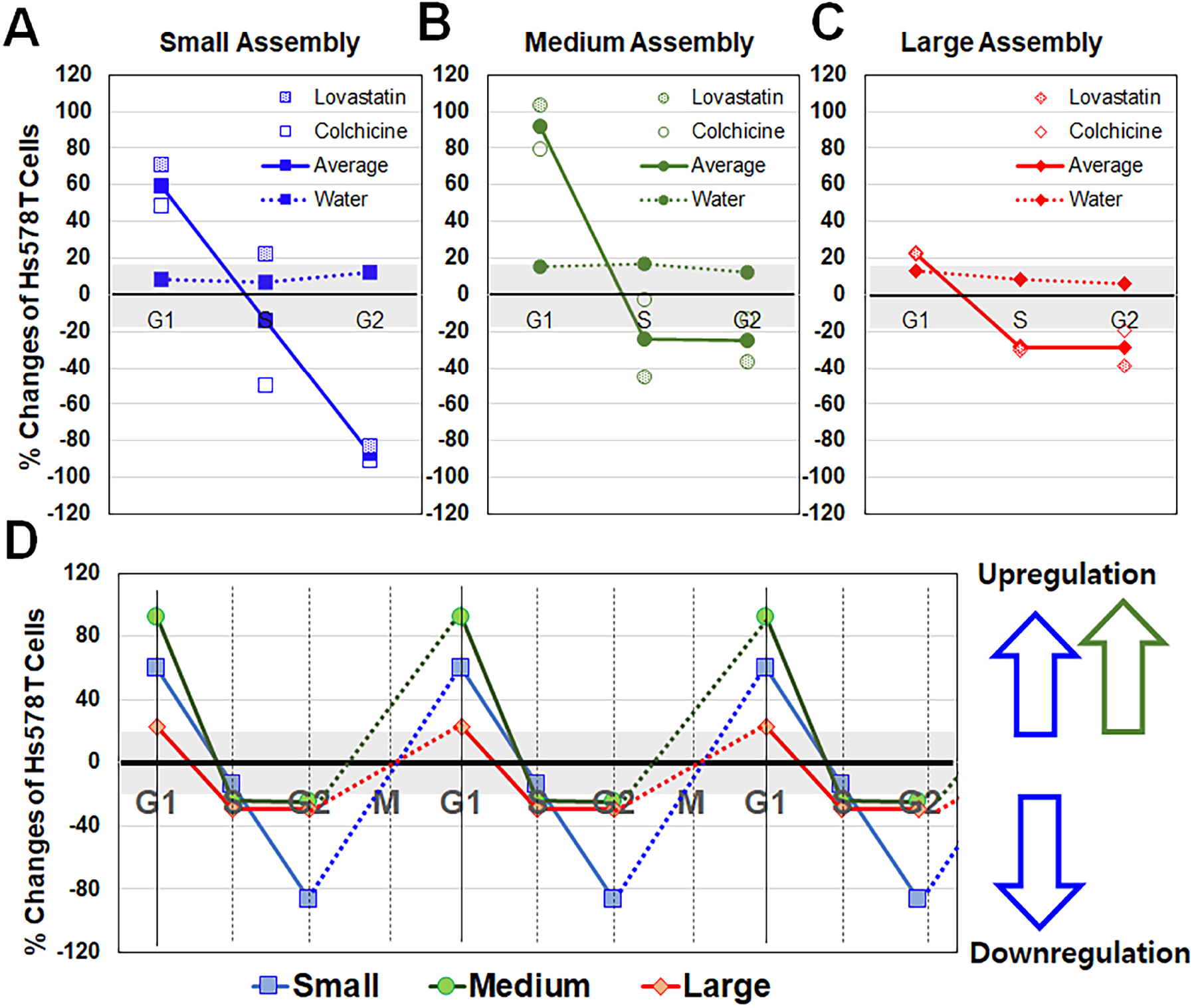
Size-dependent oscillation of glucosomes during cell cycle progression. **(A-C)** Percentage (%) changes of the number of single cells showing small- (A, blue, square), medium- (B, green, circle), and large-sized (C, red, diamond) glucosomes were analyzed along with a cell cycle (G1, S and G2) by the treatment of lovastatin (A-C, dotted symbols) and colchicine (A-C, open symbols). The average (%) changes from two cell cycle inhibitors were also analyzed along with a cell cycle (G1, S and G2) in a size-dependent manner (A-C, filled symbols with solid lines). Background changes were also observed by the treatment of a vehicle, water, as a control (A-C, filled symbols with dashed lines). **(D)** When all the data were put together in multiple cycles, size-dependent oscillation patterns of glucosomes were observed along with a cell cycle. Small-sized glucosomes (blue square) drastically oscillated during a cell cycle whereas mediumsized glucosomes (green circle) were upregulated only in the G1 phase. Large-sized glucosomes (red diamond) barely or minimally oscillated along with a cell cycle. Dashed lines indicate anticipated changes between the G2 of a cell cycle and the G1 in a next cycle. Positive and negative (%) values on y-axis indicate upregulation and downregulation of glucosomes, respectively. The grey zone (± ~17% on y-axis) indicates background noises observed by the water treatment in this analysis.

**Fig. 6.**
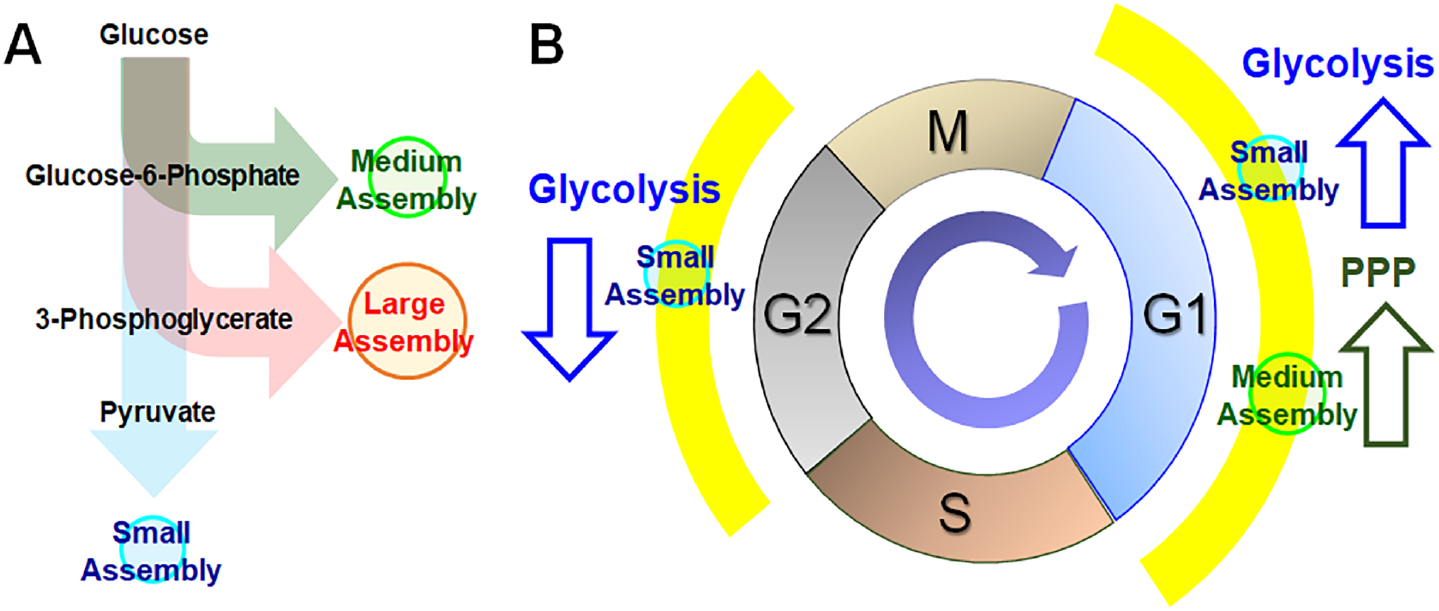
Functional analysis of glucosome dynamics during cell cycle progression. **(A)** A schematic diagram represents size-dependent functional roles of glucosomes in glucose metabolism (*15*, *19*), indicating that small-sized glucosomes primarily promote glycolysis, but medium- and large-sized glucosomes partition glucose flux preferentially into the pentose phosphate pathway and serine biosynthesis, respectively. **(B)** A schematic diagram summarizes size-dependent functional contributions of glucosomes during cell cycle progression. PPP: The pentose phosphate pathway.

## Discussion

In accordance with our previous finding of the size-dependent functional participation of glucosome assemblies in regulation of glucose flux at single-cell levels (**Fig. 6A**), small-sized glucosomes primarily promote glycolysis, but medium- and large-sized glucosomes partition glucose flux preferentially into the pentose phosphate pathway and serine biosynthesis, respectively (*15*, *19*). This means that the significantly increased numbers of Hs578T cells showing small- and medium-sized glucosomes in the G1 phase (**Figs. 4C** and **5A-B**) represent functional upregulation of glycolysis and the pentose phosphate pathway during the G1 phase (**Fig. 6B**). This is unequivocally supported by mounting amounts of literature, for example, that report an increased enzymatic activity of hexokinase 2, PFKFB3, and glucose 6-phosphate dehydrogenase, respectively, in the G1 phase and thus the upregulation of glycolysis and the pentose phosphate pathway during the G1 phase (*11*, *14*, *25*). Also, our analysis here is in line with the fact that de novo purine biosynthesis, which relies on glucose-derived carbon flux from the pentose phosphate pathway, is upregulated in the G1 phase through the formation of multienzyme purinosome assemblies (*28*). Additionally, our data showing the reduced amounts of small-sized glucosomes in the G2-arrested cells (**Figs. 4E** and **5A**) implies significant downregulation of glycolysis at the G2 phase of Hs578T cells (**Fig. 6B**). Although the activity of glycolysis is shown to diminish during the S phase relative to its level at the G1 phase (*6*, *29*), as far as we know, glycolysis has not been previously reported to be further downregulated during the G2 phase relative to its basal level activity throughout a cell cycle. Our data here represents a heretofore unidentified metabolic interplay between glycolysis and the G2 phase of a cell cycle. Furthermore, the number of Hs578T cells showing medium- and large-sized glucosomes from the S to the G2 phase (**Fig. 5B-C**) remained almost the same, indicating no alteration of glucose shunt to building block biosynthesis; in other words, a relatively sustained level of glycolysis-derived carbon flux into the pentose phosphate pathway and serine biosynthesis during the S to G2 transition. This analysis is appealing because we provide a mechanistic insight of how serine is continually generated in cancer cells to play its reported essential role in cancer cell growth and survival (*30*, *31*). Collectively, our study not only strongly supports our current understanding of metabolic demands in the G1 phase but also provides novel insights of functional interplays between glucose metabolism and a cell cycle in the G2 phase.

In addition, it is important to note here that a current model that describes temporal oscillation of metabolism during cell cycle progression has been largely established from yeast and unicellular eukaryotic cells (*6*). Particularly, metabolite profiling experiments in yeast cells demonstrated periodic changes of intracellular concentrations of metabolites, thus revealing metabolic oscillation during a cell cycle (*32*). In human cells, however, current research has been largely centered on elucidating complex mechanisms of sequential activation of cyclin-dependent kinases and their influences on metabolic enzymes. In most cases, spatial translocations of glycolytic enzymes from the cytoplasm to the nucleus (e.g., pyruvate kinase muscle-type 2, aldolase, glyceraldehyde 3-phosphate dehydrogenase, and PFKFB3) have been presented as evidence of their association with a cell cycle in human cells, rather than oscillatory alterations of their enzymatic activities in glycolysis during a cell cycle (*5*). Nevertheless, there is a report showing the activity changes of glycolytic enzymes (i.e., PFK and PKM2) by cyclin-dependent kinase 6 and thus shunting glycolytic flux into building block biosynthesis in cancer cells (*33*). In other words, we are at the infant stage of understanding whether functional oscillation of metabolic pathways occurs during a cell cycle in human cells, including cancer cells. It is also worthwhile to point out that our study provides new quantitative insights of how much glucose metabolism oscillates to meet metabolic demands during cell cycle progression in Hs578T cells (**Fig. 5**). Collectively, our study tracking glucosome dynamics from living human cells not only strengthens the importance of glucose metabolism during cell cycle progression but also reveal the behind-the-scenes mechanism of its functional oscillation during cell cycle progression through its canonical functions in metabolism.

Overall, we demonstrated that glucosomes were spatially and temporally regulated at single-cell levels and their dynamics were functionally oscillated at a population level to meet metabolic demands during cell cycle progression. Consequently, this work not only promotes the functional importance of glucosome dynamics as one of the essential metabolic entities of human cell biology, but also provides novel insights and therapeutic interventions for the treatment of human metabolic diseases, like cancer and beyond.

## Materials and Methods

### Materials

The plasmid expressing PFKL with a monomeric enhanced green fluorescent protein (PFKL-mEGFP) was previously described (*15*). Thymidine (Sigma, Cat # 89270), colchicine (EMD Millipore, Cat# 234115), and lovastatin (Tocris, Cat# 1530) were used to synchronize Hs578T cells (*23*). Vybrant^®^ DyeCycle^™^ Orange Stain (Invitrogen, Cat# V35005) was used as a cell cycle indicator.

### Cell Culture

Human triple-negative breast carcinoma Hs578T cells (HTB-126) were obtained from the ATCC. Cells were cultured in the Roswell Park Memorial Institute 1640 (RPMI 1640, Mediatech, Cat# 10-040-CV) supplemented with 10% dialyzed fetal bovine serum (dFBS) (Sigma, Cat# F2442) (*34*, *35*) and 50 μg/ml gentamicin sulfate (Corning, Cat# 61-098-RF). Cells were then maintained in a HeraCell CO_2_ incubator (37 °C, 5 % CO_2_, and 95 % humidity). The Universal Mycoplasma Detection Kit (ATCC, Cat# 30-1012K) was used to verify that cells were free of mycoplasma contamination. The Hs578T cell line was also authenticated by ATCC’s short tandem repeat profiling service.

### Transfection

Hs578T cells were gently removed from a culture flask with trypsin (Corning, Cat # 25-053-Cl) and plated on glass-bottomed 35 mm Petri dishes (MatTek) with an antibiotic-free growth medium. When cells reached ~70 - 90 % confluency, transfection was carried out using Lipofectamine 2000 (Invitrogen) with Opti-MEM-I reduced serum medium (Gibco, Cat# 11058). After 5 h of post-transfection, the Opti-MEM-I medium was exchanged with a fresh antibiotic-free growth medium that did not contain phenol red (Gibco, Cat# 11835-030) but, when desired, include a cell cycle inhibitor.

### Cell Synchronization

Hs578T cells were grown in 6-well plates ~30 - 50 % confluency in the growth medium (i.e., RPMI 1640 and 10% dFBS). On the following day, cells were transfected with PFKL-mEGFP. Cells expressing with/without PFKL-mEGFP were treated with thymidine, lovastatin, or colchicine (*22*, *23*). For the G1/S arrest, 2 mM of thymidine was added into the growth medium, and the cells were incubated for 12 h, and subsequently washed with 1x PBS twice. The growth medium was then added and incubated for 16 h. Thymidine treatment was repeated (2 mM thymidine for 12 h) before harvesting cells. For the G1 arrest, 40 μM of lovastatin was supplemented into the growth medium. After ~ 28 h incubation, cells were washed with the fresh growth medium to remove residual lovastatin. For the G2/M arrest, 100 nM colchicine was supplemented into the growth medium for 24 h. For a vehicle control, water was applied without a cell cycle inhibitor.

### Flow Cytometric Analysis

Cells were then harvested and analyzed by flow cytometry (*36*). Briefly, cells were removed from a culture plate using trypsin, and washed with 1x PBS once. Then, the cells were resuspended within ice-cold 1x PBS and fixed in ice-cold methanol for 20 min at 4 °C. The cells were washed and resuspended with 1x PBS that contains 30 μg/mL propidium iodide and 100 μg/mL RNase A (Thermo, Cat# EN0531). After incubation for at least 45 min, cells were subjected to flow cytometry using a CyAn ADP (Beckman Coulter) that is equipped with a 488 nm laser line and 530/40 and 613/20 emission filters with Summit V 4.00 software. Cell population was gated based on mEGFP intensity to exclude cells that were not expressing PFKL-mEGFP. Using the Cell Cycle platform available in FlowJo (FlowJo, LLC), cell cycle progression was determined by the Dean-Jett-Fox analysis (*37*).

### Fluorescence Live-cell Imaging

After 24 h of post-transfection of PFKL-mEGFP, Hs578T cells were washed with a buffered-saline solution (20 mM HEPES (pH 7.4), 135 mM NaCl, 5 mM KCl, 1 mM MgCl_2_, 1.8 mM CaCl_2_, and 5.6 mM glucose) for three times in a 10 min interval. To employ a cell cycle indicator, cells were subsequently treated with Vybrant^®^ DyeCycle^™^ Orange Stain (0.5 μM) for 15 min (37 °C, 5 % CO_2_, and 95 % humidity), and allowed to further incubate in the darkroom for 1–2 h at ambient temperature prior to imaging. All images were obtained using a Photometrics CoolSnap EZ monochrome CCD camera with a 60x 1.45 NA objective (Nikon CFI Plan Apo TIRF) on a Nikon Eclipse Ti inverted C2 confocal microscope. Wide-field imaging was carried out using a set of Z488/10-HC clean-up, HC TIRF dichroic, and 525/50-HC emission filter for mEGFP detection and a set of Z561/10-HC clean-up, HC TIRF dichroic, and 600/50-HC emission filter for the Vybrant^®^ DyeCycle^™^ Orange Stain detection from Chroma Technology.

### Glucosome Size Analysis

The ImageJ processing software (National Institutes of Health) was used for glucosome size analysis as reported previously (*15*). Briefly, fluorescent wide-field images were processed through ImageJ using a custom script and macro that automates the counting of fluorescent particles using its built-in module, so-called robust automatic threshold selection (RATS). In this analysis, the captured cell images were scaled according to the pixel size of the microscope (i.e., 0.12 μm/pixel) before the default parameters for RATS (i.e., noise threshold = 25, λ factor = 3) used in this analysis. Once fluorescent particles were selected from an image, the particle analysis module in ImageJ was applied to attain both the number and area of fluorescent particles within an image. This process was repeated for all subsequent cell images.

### Fluorescent Intensity Analysis from Cell Cycle Indicator

The cell cycle indicator, Vybrant^®^ DyeCycle^™^ Orange Stain, emits fluorescent signals that are proportional to the mass of DNA in single cells, which indicates cell cycle progression (*26*, *27*). To distinguish each phase of a cell cycle based on a fluorescent intensity from the nucleus, Nikon NIS-Elements software (version 4.13) was used to quantify fluorescent signals from single cells. Cells emitting the highest fluorescent intensity were served as a reference for the G2/M-arrested cells. Cells in the G1 phase were then characterized as having one-half of the reference fluorescence intensity. Any cells with a fluorescence intensity between the G1 and the G2/M phases were categorized as being in the S phase.

### Statistical Analysis

Statistical analyses of high-content imaging and flow cytometry data were performed using GraphPad Prism 9.2. Briefly, a two-way analysis of variance (ANOVA) with Tukey’s multiple comparison tests was performed to determine statistical significance among different treatments and/or between the experimental and the control groups. Statistical significance was defined as *p* < 0.05 with a 95% confidence interval.

## Acknowledgments

We thank Dr. Gregory Szeto’s group at UMBC for their assistance in operation of flow cytometry. This work is supported by the National Institutes of Health: R01GM125981 (SA), R03CA219609 (SA), R01GM134086 (MK) and T32GM066706 (MJ, DLS). The content is solely the responsibility of the authors and does not necessarily represent the official views of the National Institutes of Health.

## Author contributions

S.A. conceived the project, M.J. and D.L.S. performed experiments, and M.J., D.L.S. and M.K. carried out statistical analysis. All authors analyzed and discussed the results and wrote the manuscript.

## Data availability

All data are available within the main manuscript and the supplementary materials.

## References

1. L. Cai, B. P. Tu, Driving the cell cycle through metabolism. Annu Rev Cell Dev Biol 28 59–87 (2012).

2. I. H. Lee, T. Finkel, Metabolic regulation of the cell cycle. Curr Opin Cell Biol 25 724–729 (2013).

3. J. Kalucka, R. Missiaen, M. Georgiadou, S. Schoors, C. Lange, K. De Bock, M. Dewerchin, P. Carmeliet, Metabolic control of the cell cycle. Cell Cycle 14 3379–3388 (2015).

4. D. Roy, G. Y. Sheng, S. Herve, E. Carvalho, A. Mahanty, S. Yuan, L. Sun, Interplay between cancer cell cycle and metabolism: challenges, targets and therapeutic opportunities. Biomed Pharmacother 89 288–296 (2017).

5. P. Icard, L. Fournel, Z. Wu, M. Alifano, H. Lincet, Interconnection between metabolism and cell cycle in cancer. Trends Biochem Sci 44 490–501 (2019).

6. J. Kaplon, L. van Dam, D. Peeper, Two-way communication between the metabolic and cell cycle machineries: the molecular basis. Cell Cycle 14 2022–2032 (2015).

7. J. Chen, S. Zhang, Y. Li, Z. Tang, W. Kong, Hexokinase 2 overexpression promotes the proliferation and survival of laryngeal squamous cell carcinoma. Tumour Biol 35 3743–3753 (2014).

8. J. W. Hu, P. Sun, D. X. Zhang, W. J. Xiong, J. Mi, Hexokinase 2 regulates G1/S checkpoint through CDK2 in cancer-associated fibroblasts. Cell Signal 26 2210–2216 (2014).

9. S. Duan, M. Pagano, Linking metabolism and cell cycle progression via the APC/CCdh1 and SCFbetaTrCP ubiquitin ligases. Proc Natl Acad Sci U S A 108 20857–20858 (2011).

10. W. Jia, X. Zhao, L. Zhao, H. Yan, J. Li, H. Yang, G. Huang, J. Liu, Non-canonical roles of PFKFB3 in regulation of cell cycle through binding to CDK4. Oncogene 37 1685–1698 (2018).

11. F. Peng, Q. Li, J. Y. Sun, Y. Luo, M. Chen, Y. Bao, PFKFB3 is involved in breast cancer proliferation, migration, invasion and angiogenesis. Int J Oncol 52 945–954 (2018).

12. L. Shi, H. Pan, Z. Liu, J. Xie, W. Han, Roles of PFKFB3 in cancer. Signal Transduct Target Ther 2, 17044 (2017).

13. A. Benito, I. H. Polat, V. Noe, C. J. Ciudad, S. Marin, M. Cascante, Glucose-6-phosphate dehydrogenase and transketolase modulate breast cancer cell metabolic reprogramming and correlate with poor patient outcome. Oncotarget 8, 106693–106706 (2017).

14. P. Vizan, G. Alcarraz-Vizan, S. Diaz-Moralli, O. N. Solovjeva, W. M. Frederiks, M. Cascante, Modulation of pentose phosphate pathway during cell cycle progression in human colon adenocarcinoma cell line HT29. Int J Cancer 124, 2789–2796 (2009).

15. C. L. Kohnhorst, M. Kyoung, M. Jeon, D. L. Schmitt, E. L. Kennedy, J. Ramirez, S. M. Bracey, B. T. Luu, S. J. Russell, S. An, Identification of a multienzyme complex for glucose metabolism in living cells. J Biol Chem 292, 9191–9203 (2017).

16. M. Kyoung, S. J. Russell, C. L. Kohnhorst, N. N. Esemoto, S. An, Dynamic architecture of the purinosome involved in human de novo purine biosynthesis. Biochemistry 54, 870–880 (2015).

17. M. Jin, G. G. Fuller, T. Han, Y. Yao, A. F Alessi, M. A. Freeberg, N. P. Roach, J. J. Moresco, A. Karnovsky, M. Baba, J. R. Yates, 3rd, A. D. Gitler, K. Inoki, D. J. Klionsky, J. K. Kim, Glycolytic Enzymes Coalesce in G Bodies under Hypoxic Stress. Cell reports 20, 895–908 (2017).

18. B. A. Webb, A. M. Dosey, T. Wittmann, J. M. Kollman, D. L. Barber, The glycolytic enzyme phosphofructokinase-1 assembles into filaments. The Journal of cell biology 216, 2305–2313 (2017).

19. M. Jeon, H. W. Kang, S. An, A mathematical model for enzyme clustering in glucose metabolism. Sci Rep 8, 2696 (2018).

20. A. M. Abukhdeir, B. H. Park, P21 and p27: roles in carcinogenesis and drug resistance. Expert Rev Mol Med 10, e19 (2008).

21. J. Laurent, C. Frongia, M. Cazales, O. Mondesert, B. Ducommun, V. Lobjois, Multicellular tumor spheroid models to explore cell cycle checkpoints in 3D. BMC cancer 13 73 (2013).

22. G. Chen, X. Deng, Cell synchronization by double thymidine block. Bio Protoc 8, (2018).

23. J. Jackman, P. M. O’Connor, Methods for synchronizing cells at specific stages of the cell cycle. Curr Protoc Cell Biol Chapter 8, Unit 8 3 (2001).

24. A. L. Blajeski, V. A. Phan, T. J. Kottke, S. H. Kaufmann, G(1) and G(2) cell-cycle arrest following microtubule depolymerization in human breast cancer cells. J Clin Invest 110, 91–99 (2002).

25. L. Li, S. Jiang, X. Li, Y. Liu, J. Su, J. Chen, Recent advances in trimethoxyphenyl (TMP) based tubulin inhibitors targeting the colchicine binding site. Eur J Med Chem 151, 482–494 (2018).

26. K. Bedelbaeva, A. Snyder, D. Gourevitch, L. Clark, X. M. Zhang, J. Leferovich, J. M. Cheverud, P. Lieberman, E. Heber-Katz, Lack of p21 expression links cell cycle control and appendage regeneration in mice. Proc Natl Acad Sci U S A 107, 5845–5850 (2010).

27. D. T. Dicker, J. M. Lerner, W. S. El-Deiry, Hyperspectral image analysis of live cells in various cell cycle stages. Cell Cycle 6, 2563–2570 (2007).

28. C. Y. Chan, H. Zhao, R. J. Pugh, A. M. Pedley, J. French, S. A. Jones, X. Zhuang, H. Jinnah, T. J. Huang, S. J. Benkovic, Purinosome formation as a function of the cell cycle. Proc Natl Acad Sci U S A 112 1368–1373 (2015).

29. Y. Bao, K. Mukai, T. Hishiki, A. Kubo, M. Ohmura, Y. Sugiura, T. Matsuura, Y. Nagahata, N. Hayakawa, T. Yamamoto, R. Fukuda, H. Saya, M. Suematsu, Y. A. Minamishima, Energy management by enhanced glycolysis in G1-phase in human colon cancer cells in vitro and in vivo. Mol Cancer Res 11, 973–985 (2013).

30. B. Chaneton, P. Hillmann, L. Zheng, A. C. L. Martin, O. D. K. Maddocks, A. Chokkathukalam, J. E. Coyle, A. Jankevics, F. P. Holding, K. H. Vousden, C. Frezza, M. O’Reilly, E. Gottlieb, Serine is a natural ligand and allosteric activator of pyruvate kinase M2. Nature 491, 458–462 (2012).

31. O. D. Maddocks, C. R. Berkers, S. M. Mason, L. Zheng, K. Blyth, E. Gottlieb, K. H. Vousden, Serine starvation induces stress and p53-dependent metabolic remodelling in cancer cells. Nature 493, 542–546 (2013).

32. B. P. Tu, R. E. Mohler, J. C. Liu, K. M. Dombek, E. T. Young, R. E. Synovec, S. L. McKnight, Cyclic changes in metabolic state during the life of a yeast cell. Proc Natl Acad Sci USA 104, 16886–16891 (2007).

33. H. Wang, B. N. Nicolay, J. M. Chick, X. Gao, Y. Geng, H. Ren, H. Gao, G. Yang, J. A. Williams, J. M. Suski, M. A. Keibler, E. Sicinska, U. Gerdemann, W. N. Haining, T. M. Roberts, K. Polyak, S. P. Gygi, N. J. Dyson, P. Sicinski, The metabolic function of cyclin D3-CDK6 kinase in cancer cell survival. Nature 546, 426–430 (2017).

34. S. An, M. Jeon, E. L. Kennedy, M. Kyoung, Phase-separated condensates of metabolic complexes in living cells: purinosome and glucosome. Methods Enzymol 628, 1–17 (2019).

35. S. An, R. Kumar, E. D. Sheets, S. J. Benkovic, Reversible compartmentalization of de novo purine biosynthetic complexes in living cells. Science 320, 103–106 (2008).

36. L. C. Crowley, G. Chojnowski, N. J. Waterhouse, Measuring the DNA content of cells in apoptosis and at different cell-cycle stages by propidium iodide staining and flow cytometry. Cold Spring Harb Protoc 2016, (2016).

37. M. H. Fox, A model for the computer analysis of synchronous DNA distributions obtained by flow cytometry. Cytometry 1, 71–77 (1980).

