## Supplementary Materials for "Functional oscillation of a multienzyme glucosome assembly during cell cycle progression"

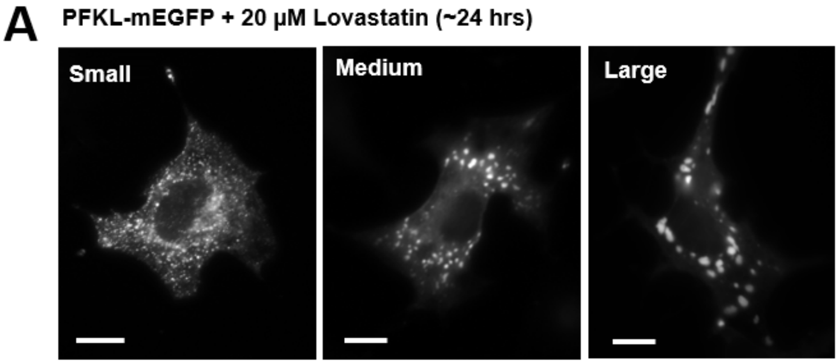


**B**

| % Cells showing differently sized glucosomes | | | |
|---|---|---|---|
| | Small | Medium | Large |
| Hs578T (PFKL-mEGFP) | **58.8** ± 4.9% | **13.3** ± 3.2% | **26.5** ± 3.6% |
| + 20 µM Lovastatin | **66.9** ± 5.0% | **21.1** ± 6.1% | **12.0** ± 6.0% |

**Supplementary Fig. S1. Effect of lovastatin on cell population showing differently sized glucosomes.** The population (%) of Hs578T cells displaying each size of PFKL-mEGFP assemblies was analyzed in the presence of 20 µM lovastatin ($N_{lovastatin}$ = 5). **(A)** Representative images were obtained from lovastatin-treated Hs578T cells displaying three different sizes of glucosomes. Scale bars,10 µm. **(B)** A table summarizes changes of the numerical values of the averaged percentages (%) of cells displaying given sized glucosomes in the absence and presence of lovastatin. Error ranges in (B) represent standard deviations from at least five independent trials.

A PFKL-mEGFP + 10 nM Colchicine (~24 hrs)

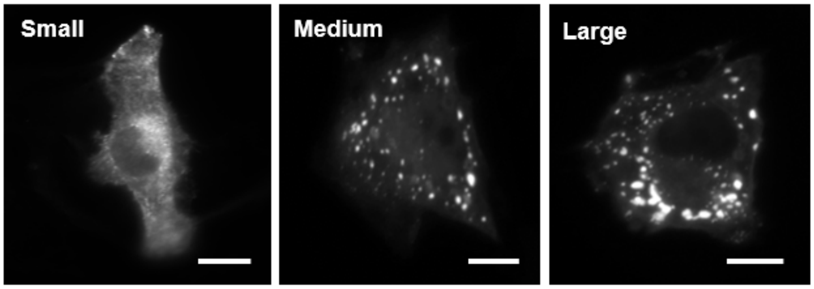


B

| % Cells showing differently sized glucosomes | | | |
|---|---|---|---|
| | Small | Medium | Large |
| Hs578T (PFKL-mEGFP) | **58.8** ± 4.9% | **13.3** ± 3.2% | **26.5** ± 3.6% |
| + Colchicine (10 nM) | **75.1** ± 8.6% | **11.4** ± 3.2% | **13.6** ± 5.6% |

**Supplementary Fig. S2. Effect of colchicine on cell population showing differently sized glucosomes.** The population (%) of Hs578T cells displaying each size of PFKL-mEGFP assemblies was analyzed in the presence of 10 nM colchicine ($N_{colchicine} = 5$). **(A)** Representative images were obtained from colchicine-treated Hs578T cells displaying three different sizes of glucosomes. Scale bars,10 μm. **(B)** A table summarizes changes of the numerical values of the averaged percentages (%) of cells displaying given sized glucosomes in the absence and presence of colchicine. Error ranges in (B) represent standard deviations from at least five independent trials.
